# Electrophysiological and network dynamics disruption of hippocampal CA1 neurons after NMDAr blockade by MK-801

**DOI:** 10.1101/2025.01.28.635254

**Authors:** P. Abad-Perez, F. J. Molina-Payá, G. Rigamonti, M Tirado, G. Cabrales, Antonio Falcó Montesinos, L. Martínez-Otero, V. Borrell, J. Sánchez-Mut, R. Redondo, J. R. Brotons-Mas

**Affiliations:** Universidad Cardenal Herrera-CEU, CEU Universities, Spain; Instituto de Neurociencias UMH-CSIC, Alicante, Spain; IMIM, Hospital del Mar Research Institute; Roche Pharma Research and Early Development, Neuroscience and Rare Diseases, Roche Innovation Center Basel, F. Hoffmann-La Roche Ltd, Grenzacherstrasse 124, 4070 Basel, Switzerland

**Keywords:** Oscillations, single neuron, NMDAr, schizophrenia

## Abstract

The N-methyl-D-aspartate receptor (NMDAr) hypofunction hypothesis suggests that excitatory-inhibitory (E-I) imbalance underlies cognitive deficits associated with neuropsychiatric disorders, such as schizophrenia (SCZ). In this study, we investigated the impact of NMDAr blockade on pyramidal neurons and interneurons in the CA1 region of the hippocampus using in vivo electrophysiological recordings during both spontaneous and exploratory behaviors. Through the administration of MK-801, an NMDAr antagonist, we assessed changes in spike train dynamics, network synchrony, and neuronal modulation by oscillatory activity, with particular emphasis on theta, gamma, and sharp-wave ripple (SWR) oscillations. We found that NMDAr blockade significantly disrupted the E-I balance, leading to altered spike train properties, reduced bursting propensity, and impaired neuronal synchronization. These changes were accompanied by decreased modulation of pyramidal neurons and interneurons by theta and gamma oscillations, as well as diminished recruitment of pyramidal neurons during SWR events. Additionally, the correlation between firing rates and movement speed was reduced, reflecting deficits in spatial coding and memory processing. Moreover, MK-801 administration disrupted place cell stability and spatial information processing. These effects likely contribute to functional disconnection between the hippocampus and other brain regions, such as the prefrontal cortex, and may underlie SCZ-associated cognitive impairments. Our findings provide valuable insights into the cellular and network-level mechanisms affected by NMDAr dysfunction, highlighting the role of oscillatory activity and spike timing in cognitive deficits. This work advances our understanding of how NMDAr hypofunction impacts hippocampal circuits and identifies potential targets for therapeutic interventions aimed at restoring cognitive function in neuropsychiatric disorders.

## Introduction

A major challenge in the field of neuroscience is to understand how neuronal activity sustains complex mental processes in physiological and pathological conditions. In this sense, it is believed that brain rhythms, the result of the interaction between pyramidal excitatory neurons and inhibitory interneurons, are one of the key mechanisms underlying neuronal computations (Buzsáki & Watson, 2012). Given this, it is not surprising that the presence of aberrant oscillatory activity in different neuropsychiatric illnesses(Uhlhaas & Singer, 2010). One example is schizophrenia (SCZ) a disorder which manifests by negative, positive, and cognitive symptoms (Owen et al., 2016) and that is characterize by aberrant gamma (30-100Hz) oscillations and parvalbumin positive interneurons (PV+) disfunction (Dienel & Lewis, 2019; Uhlhaas & Singer, 2010)

Although there have been multiple efforts to explain the origin of SCZ, its etiology remains unknow ((Ban, 2007; Lewis & Lieberman, 2000; Meltzer & Stahl, 1976). However, it is generally assumed that SCZ emerges as result of the interaction between genetic factors, viral infection, psychosocial stress during critical neurodevelopmental periods (Insel, 2010). The confluence of these influences would alter the normal activity of interneurons and brain circuits generating abnormal oscillatory activity (Del Pino et al., 2013; Hobson et al., 2021; Laurent et al., 2015; O’Neill et al., 2013; Sigurdsson et al., 2010).

The NMDAr hypofunction hypothesis aims to explain SCZ as a consequence of interneuron dysfunction. Specifically, the loss of NMDAr function alters the excitation-inhibition (E-I) balance, inducing aberrant oscillatory activity and SCZ symptoms (Lee & Zhou, 2019). In fact, the administration of NMDAr antagonists can induce SCZ-like symptoms in both patients and healthy subjects (Adler et al., 1998, 1999; Malhotra et al., 1996). Similarly, in animal models, NMDAr blockade using different receptor antagonists (dizocilpine/MK-801, phencyclidine/PCP, or ketamine) recapitulates many behavioral and cognitive symptoms observed in SCZ. Moreover, these treatments induce aberrant oscillatory activity resembling the behavioral and cognitive alterations observed in patients (Corbett et al., 1995; Ma and Leung, 2007; Chrobak et al., 2008; Pitsikas et al., 2008; Kubík et al., 2014; Cui et al., 2022; (Abad-Perez et al., 2023; Chrobak et al., 2008; Corbett et al., 1995; Cui et al., 2022; Kubík et al., 2014; Ma & Leung, 2007; Maleninska et al., 2022; Pitsikas et al., 2008). Additionally, the deletion of NMDAr from PV+ interneurons in mice also induces SCZ-like symptoms (Lee & Zhou, 2019; Ohtsuki et al., 2001).

Single-neuron activity analyses after MK-801 administration reveal a disruption in the balance of excitatory and inhibitory neuronal activity recorded in the PFC. This results in disorganized neuronal activity (Homayoun & Moghaddam, 2007; Jackson et al., 2004; Molina et al., 2014) and a progressive shift in the firing rates of excitatory and inhibitory interneurons (Jackson et al., 2004; Homayoun and Moghaddam, 2007; Molina et al., 2014). In the hippocampus, NMDAr blockade disrupts single-neuron activity, spatial representation stabilization during goal navigation tasks, and memory refinement during consolidation (Dupret et al., 2010; Kentros et al., 1998; Norimoto et al., n.d.).

In our previous work (Abad-Perez et al., 2023), we demonstrated that NMDAr blockade disrupts the normal relationship between oscillatory activity and behavior. Furthermore, we observed impaired dialog between the hippocampus and the prefrontal cortex (PFC), marked by disrupted theta/gamma co-modulation during a spatial working memory task. Additionally, we observed specific effects of MK-801 in the hippocampus and PFC. We hypothesized that these changes, especially in theta and gamma co-modulation activity, could be linked to alterations in neuronal function, including changes in firing rates, spike train organization, and the loss of network/neuron modulation.

In this work, we aimed to investigate the effects of NMDAr blockade on pyramidal excitatory neurons and narrow-waveform inhibitory interneurons recorded in the CA1 region of the hippocampus. We also expected to observe changes in unit activity during ripples, as well as alterations in network synchrony and bursting activity. Alterations in these parameters might explain deficits in spatial information processing and help account for the functional disconnection between the hippocampus and the PFC. Our results indicate that NMDAr blockade induces many of these effects.

## Methods

### Subjects

Male wild-type mice C57Bl6/J, N=6 aged p60 to p90 supplied by the “RMG” were used for the study. Mice were maintained on a 12 h light/dark cycle with food and water available ad libitum and were individually housed in standard cages after electrode implantation. All experimental procedures were approved by the UMH-CSIC ethics committee and the regional government and met local and European guidelines for animal experimentation (86/609/EEC).

### In vivo recordings on freely moving mice

Microdrives, (Axona ltd) mounting four independently movable tetrodes (12 μm tungsten wire, California Fine Wire Company, Grover Beach, CA, USA) or 3D-printed resin microdrives (Vandecasteele et al., 2012) mounting silicon probes (NeuroNexus Tech, Ann Arbor, MI, USA), were implanted under isoflurane anesthesia (1.5%) while buprenorphine (0.05mg/kg, s.c.) was supplied as analgesia. Craniotomies were performed above the hippocampus, targeting the hippocampal CA1 region (AP: −81.8-2.2, M-L: 1.2 V: 0.6 mm). Animals were left to recover for at least seven days after surgery (Brotons-Mas et al., 2010, 2017; Del Pino et al., 2017).

### Recordings and behavior

Electrophysiological data were acquired using an Intan base system (NeuroNexus Tech, Ann Arbor, MI, USA) digitized with 30 kHz rate. The wide-band signal was downsampled to 1.25 kHz and used as the LFP signal. The animal’s position was recorded using Any-Maze (Stolteing, LTD) sampling at 30Hz. Position was synchronized with neural data with TTLs signaling. Animals were handled daily and accommodated to the experimenter, recording room and cables for 1 week before the start of the experiments. Mice were recorded during spontaneous exploration in an open field (50×50cm) and in their home cages.

### Behavioral Protocol and drug administration

Once animals recovered from surgeries, electrodes aiming for CA1 were lowered. Unit activity, ripples and theta power were used as electrophysiological landmarks to determine electrode’s location in the CA1 pyramidal layer. Experiments began with a baseline condition. This served to determine change due to drug administration controlling for possible electrode position drifting across days.

Each baseline and drug condition consisted of several phases, see (Abad-Perez et al., 2023). First, recordings were performed in the animal’s home cages for 1h (HC1). Then, electrical activity was recorded during spontaneous exploration in an open field of size 50 x 50 cm (OF1) for 20 minutes. Animals were then placed again in their home cages and were left to rest for 1h (HC2). Subsequently, mice were injected subcutaneously wither with MK-801 (0.075 mg/kg) diluted in 0.3% tween80 in saline or with the vehicle solution, consisting of diluted in 0.3% tween80 in saline (Abad et al, 2023) and recordings(Petersen et al., n.d.) were performed in the home cage for 20 min (HC3). 20 minutes after the injection, mice were placed again in the open field for spontaneous exploration for 15 minutes (OF’) within the temporal window of MK-801 bioavailability, maximum concentration 40-60 min after administration and slowly decaying up to 120 min later (Wegener et al, 2011; Abad et al, 2023). Finally, animals were left to rest in their home cages and recordings were performed for one hour (HC4).

Drugs were administered following a counterbalanced scheme across alternate days. This way each mouse either received MK-801 or vehicle injections on day one, and the complementary treatment on day two. Experiments were spaced at least 48h.

### Spike sorting and neuron classification

Spike sorting was performed semi-automatically with KiloSort (Pachitariu et al, 2016; https://github.com/cortex-lab/KiloSort), using the KilosortWrapper pipeline (a wrapper for KiloSort, https://github.com/brendonw1/KilosortWrapper). This was followed by manual adjustment of the waveform clusters using the software Phy (https://github.com/kwikteam/phy) and specific plugins for phy (https://github.com/petersenpeter/phy-plugins). The following parameters were used for the Kilosort clustering: ops.Nfilt: 6 * numberChannels; ops.nt0: 64; ops.whitening: ‘full’; ops.nSkipCov: 1; ops.whiteningRange: 64; ops.criterionNoiseChannels: 0.00001; ops.Nrank: 3; ops.nfullpasses: 6; ops.maxFR: 20000; ops.fshigh: 300; ops.ntbuff: 64; ops.scaleproc: 200; ops.Th: [4 10 10]; ops.lam: [5 20 20]; ops.nannealpasses: 4; ops.momentum: 1./[20 800]; ops.shuffle_clusters: 1. Units were separated into putative pyramidal cells and narrow waveform interneurons using their autocorrelograms, waveform characteristics and firing rate. The classification was performed using CellExplorer (https://cellexplorer.org/; (Petersen et al., n.d.)).

### Power spectrum analysis

Local field potentials were analyzed using custom-written Matlab codes (Mathworks, Natick, MA, USA). Spectral power was calculated using the Thomson multi-taper method included in Chronux signal processing toolbox (Bokil et al., 2010).

### Ripple detection

Ripples were detected as described previously described (Tingley and Buzsáki, 2020), and the code (findRipples) is available online (https://github.com/valegarman/HippoCookBook). The raw LFP (1250 Hz) was filtered (130-200 Hz; Butterworth; order = 3) and transformed to a normalized squared signal (NSS). This signal was used to identify peaks that were beyond 5 SD above the mean NSS. The beginning/end of the ripple was defined by a threshold of 2 SD above the mean NSS. Ripple duration limits were between 30ms and 100ms. In addition, estimated EMG from the LFP ((Schomburg et al., 2014)) was used to exclude EMG related artifacts. The peak of the ripple (max power value > 5 standard deviations) was defined as time 0 for the ripple. Ripples were defined across the whole session recording. Then, ripples were divided into baseline and drug conditions. The channel used to identify ripples was selected manually to improve the detection. The sample ripple channel was used for computing phase modulation of different frequency bands (ripple, theta, low gamma, and high gamma).

### Ripples response

To compute ripple recruitment of single units we calculated ripple triggered peri-events histograms (PETHs) by counting spiking activity around peak ripple time into 5 ms bins. The response curve was then smoothed using a 5-point moving average fulter (‘smooth’ function in Curve Fitting Toolbox ^TM^ in Matlab). A Z-scored response curve was computed by substracting and dividing by the standard deviation of the response curve in the −500 ms to −25 ms time window. Firing rate during ripple events were defined by computing the mean firing rate in a −25 ms to 25 ms window around ripple peak. Baseline firing rate was defined by computing the mean firing rate in a −500 ms to −450 ms time window with respect to ripple peaks. Units were deemed significantly SPW-R modulated using three distinct criteria. First, a bootstrap test was computed. 500 surrogate PETHs were created for each unit by selecting random timestamps (same number as ripple events) between the first and last ripple peak event. Mean and confidence intervals of the firing rate during random points were calculated. When the mean firing rate during ripple events exceeded the biggest confidence interval value for random events, the units was considered significantly positive modulated by ripple events. Second, a two-sample Kolmogorov-Smirnov test (‘kstest2’ function in Statistics Toolbox in Matlab) was used to compare the firing rate during ripple events to the baseline firing rates. We defined modulated units by these criteria only when the associated p-value of the test was lower than 0.01. Third, if the Z-scored firing rate during ripple peak events was bigger than 1.96, the unit was considered significantly positive modulated by ripple events. Only when all three criteria were significant, units were defined as significantly ripple modulated. Only units significantly modulated during the baseline condition were used for both the ripple response and phase modulation analysis.

### Phase locking to ongoing oscillations

Phase locking of single units to ongoing oscillations was computed as previously described (Zutshi et al 2022) and the code (phaseModulation) is available online (https://github.com/valegarman/HippoCookBook). The LFP was filtered for each frequency of interest and the Hilbert transform was applied to obtain the instantaneous phase. Phase locking was computed by calculating the phase angles for each timestamp where an action potential occurred. A histogram of phase angles was calculated, and the circular mean and resultant vector (Berens et al 2009) were calculated for each neuron. Only neurons that were significantly modulated (Rayleigh test) for each frequency band during the baseline condition were included for the analysis. Ripple modulation was not the case, since only significant responsive neurons for ripple events during the baseline condition were included both in the analysis of responsiveness and phase modulation.

### Cross-frequency coupling modulation index

A cross-frequency modulation index (MI) was computed based on a normalized entropy measure as previously described (Tort et al 2008). The raw LFP signal was filtered in the theta band (4-12 Hz) and in different frequency intervals for gamma band: low gamma (20-60 Hz) and high gamma (60-100 Hz). Then, the Hilbert transform of the filtered signals were computed and the phase of theta oscillations and the envelope of gamma were obtained. Theta phases were binned into eighteen 20° intervals (0° to 360°) and the mean amplitude of all frequency bands for each phase was calculated. Finally, we applied the entropy measure, defined by:

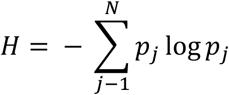

Where N = 18 (number of bins) and *p_j_* represents the mean amplitude for each binned phase *j*. The MI is finally obtained by normalizing *H* by the maximum possible entropy value *H_max_*, which is obtained for the uniform distribution 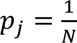 (and hence *H_max_* = *logN*):

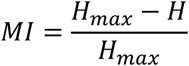

A MI value of 0 indicates lack of phase-to-aplitude modulation (constant amplitude for all phase bins), and larger MI values result from stronger phase-to-amplitude modulation.

### Firing rate and speed of movement correlation

We computed the instantaneous firing rate of CA1 neurons in each position sample (3 ms bin) and computed the Pearson correlation between the instantaneous firing rate and the speed of movement, obtaining a speed score correlation coefficient and an associated p-value. Epochs of quietness were not included in the analysis. Only significant positive-correlated cells for both baseline and drug condition were analyzed

### Anatomical verification of electrode location

After completing experiments, animals were sacrificed using sodium pentobarbital (180mg/kg at a concentration of 100mg/ml) and transcardially perfused with saline and 4% paraformaldehyde. This procedure was authorized by the ethical commission according to the current normative and following veterinary recommendations. The brains were removed, sliced in 40 µm sections, and prepared for Immunohistochemistry against DAPI, NeuN (Merck,ltd) and GFAP (Adigene technologies) for electrode localization.

### Statistical analysis

Variables distribution was tested for normality using Kolmogorov-Smirnov test. To determine statistically significant differences across conditions, parametric and non-parametric tests were used. ANOVA repeated measures was used for normally distributed variables The Friedman test was used for non-normally distributed variables. Then, paired comparisons between specific conditions were tested using t-test for repeated measures and non-parametric Wilcoxon test when appropriate. In the case of correlations, normality was verified, and the parametric Pearson coefficient were calculated. All descriptive values were expressed as mean ± the standard error (S.E.). Statistical significance was set at p< 0.05, and Bonferroni corrections were used when necessary. All statistical analysis was performed using matlab statistic toolbox.

## Results

### NMDAr blockade by MK-801 altered spike train dynamics

Previous studies indicate that blocking NMDAr alters oscillatory activity, leading to rhythms and cognitive deficits akin to those observed in SCZ. At the single-neuron level, NMDAr blockade disrupts neuronal activity in the PFC, significantly impacting the excitatory-inhibitory balance (Homayoun & Moghaddam, 2007; Jackson et al., 2004; Molina et al., 2014). Moreover, the impact of NMDAr blockade on the oscillatory profile varies across regions (Abad-Perez et al., 2023).

The hippocampus contains very complex circuits crucial for memory processing and supporting different rhythms, such as theta, gamma and sharp-wave ripples (SWR). These rhythms heavily rely on the proper excitatory-inhibitory balance (E-I), which are strongly affected in SCZ. Thus, we investigated the presence of a specific profile of activity in the CA1 region after MK-801 administration. We performed LFP recordings and signal processing to investigate hippocampal rhythms and single neuron activity performing spike sorting, and then we classified neurons as putative pyramidal (PYR) and interneuron (INT) (Petersen et al 2020; Brotons-Mas et al, 2017).

A total of N = 358 units were recorded across the baseline-vehicle condition, and N = 465 units were recorded during the baseline-MK-801 experiments. In the baseline-vehicle condition, n = 289 were identified as pyramidal neurons (PYR), and n = 69 were classified as interneurons (INT) (Fig. 1, A–C). Similarly, in the baseline-MK-801 condition, n = 400 units were classified as pyramidal neurons, and n = 65 were identified as interneurons. We investigated the general electrophysiological properties of PYR and INT. No significant changes in the firing rates of PYR and INT were observed after treatment with MK-801 (Baseline PYR: 1.48 ± 0.09 Hz; MK-801 PYR: 1.54 ± 0.09 Hz; p = 0.13; Baseline INT: 9.33 ± 1.27 Hz; MK-801 INT: 9.5 ± 1.4 Hz; p = 0.17). In contrast, the firing activity was significantly diminished in the vehicle condition for both PYR and INT (Baseline PYR: 1.78 ± 0.1 Hz; Vehicle PYR: 1.66 ± 0.08 Hz; p < 0.001; Baseline INT: 6.96 ± 1.34 Hz; Vehicle INT: 6.77 ± 1.23 Hz; p < 0.001) (Fig. 1).

**Figure 1:**
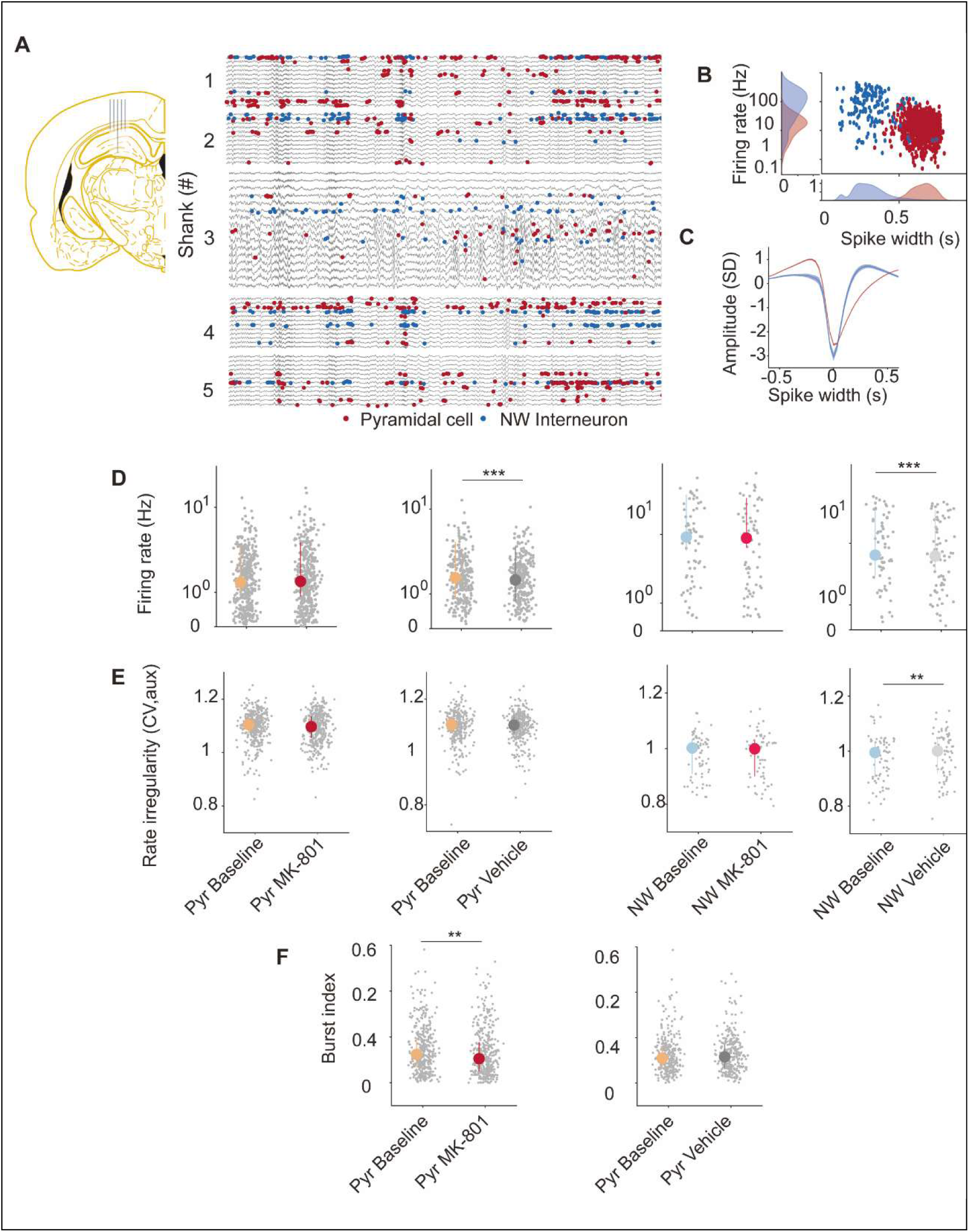
A. Raw recording and B. Spike waveform and firing rate were used to classify neurons as excitatory or inhibitory. Note the two cluster in B. C and D. Pyramidal neurons augmented their firing rate after NMDAr blockade, while diminished in the vehicle condition. E and F. Pyramidal burstiness was reduced after NMDAr blockade. G. and H. Interneurons, followed a similar profile to that of PYR, but only reached significant values for the reduction in the firing rate during the vehicle condition.

Further, we hypothesized that NMDAr blockade should induce changes in the spike train dynamics of pyramidal neurons (PYR). To test this, we investigated the coefficient of firing rate variability of PYR and interneurons (INT). We found no significant differences except for the vehicle condition that was somehow increased for interneurons IN in the vehicle condition (BS PYR MK-801: 1.1 ± 0.003, MK-801: 1.1 ± 0.003, p = 0.13; BS INT MK-801: 1 ± 0.01, MK-801: 1 ± 0.01, p = 0.71; BS PYR Vehicle: 1.1 ± 0.003, Vehicle: 1.1 ± 0.003, p = 0.37; BS INT Vehicle: 0.99 ± 0.01, INT Vehicle: 1.1 ± 0.01 p>0.01). Next, we examined the bursting properties of PYR and INT (Mizuseki et al., 2012; Petersen et al., 2020). Our analysis revealed a significant decrease in the bursting propensity of PYR following NMDAr blockade, whereas no significant differences were observed in the vehicle condition (Figure 1; Baseline MK-801: 0.12 ±0.005; MK-801: 0.1 ± 0.005; p = 0.01; Baseline Vehicle: 0.11 ± 0.005; Vehicle: 0.11 ± 0.005; p = 0.26).

Finally, we investigated population synchrony by calculating the population cross-correlogram. This type of analysis is relevant to determine the degree of circuit coordination. Our analysis revealed that MK-801 administration resulted in a reduction in the synchrony in PYR and INT, whereas an opposite trend was observed in the vehicle condition (Figure 2; PYR Baseline MK-801: 1.37± 0.03; PYR MK-801: 1.13 ± 0.03; p < 0.001; INT BS MK-801: 1.58 ± 0.07, INT MK-801: 1.49 ± 0.06, p = 0.02; PYR BS Vehicle: 1.23 ± 0.03, PYR Vehicle: 1.37 ± 0.03, p < 0.001; INT BS Vehicle: 1.64 ± 0.08, INT Vehicle: 1.76 ± 0.07, p = 0.002).

**Figure 2:**
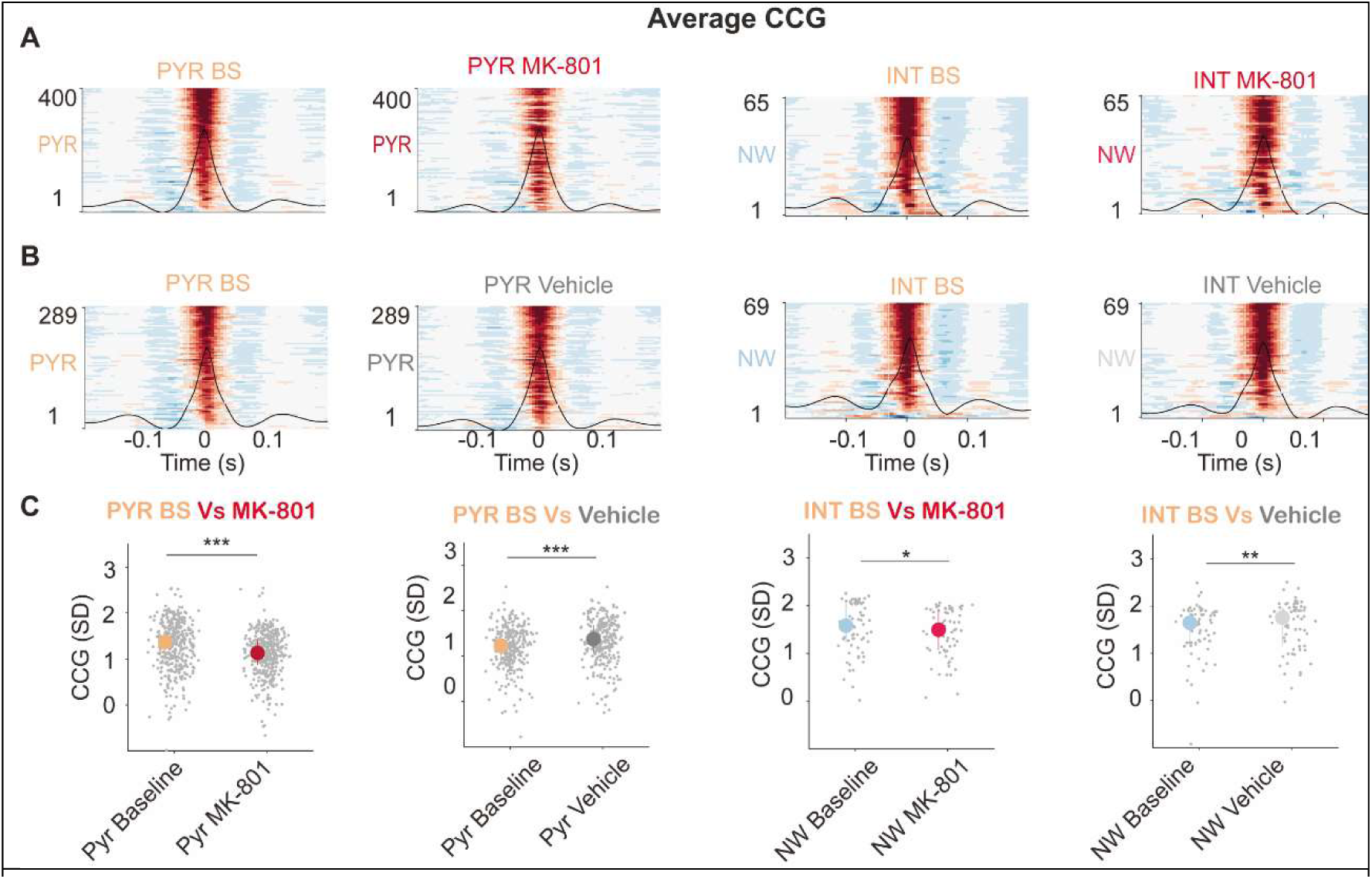
From left to right, heat maps represent single-neuron activity with respect to simultaneously recorded neurons. A. Cross-correlogram for PYR and INT under baseline and MK-801 conditions. Note the reduced firing synchrony of PYR neurons in the MK-801 condition. B. Cross-correlogram for PYR and INT under baseline and vehicle conditions. C. Quantification of the data shown in A and B. PYR firing synchrony significantly decreased in the MK-801 condition compared to baseline, whereas an increase in synchrony was observed in the vehicle condition.

Therefore, NMDAr blockade did not change the firing rate across the population of neurons recorded but distorted the spike train properties and the coordination of the neuronal network.

### NMDAr blockade altered the modulation of single neuron activity by the ongoing oscillations

NMDAr blockers can significantly alter the oscillatory profile, disrupting communication between different brain regions (Abad-Perez et al., 2023; Bygrave et al., 2019). In this study, we observed similar alterations in oscillatory activity following NMDAr blockade (data not showed). Furthermore, we aimed to investigate whether these oscillatory alterations corresponded to network-level changes in single-neuron activity.

First, we investigated the relationship between pyramidal neurons (PYR), interneurons (INT), and theta oscillations. The fidelity of this association serves as a reliable biomarker of physiological function and as an indicator of information processing. Our analysis revealed that NMDAr blockade significantly advanced the preferred firing phase of pyramidal neurons (PYR) toward earlier phases of theta cycles and reduced the strength of phase-locking. This reduction in phase-locking strength was also observed in interneurons (INT) (see Fig. XXX) (BS MK-801 PYR Phase: 4 ± 0.04 rad; MK-801 PYR Phase: 3.76 ± 0.05 rad; *p* = 0.002; BS MK-801 PYR MVL: 0.26 ± 0.01; MK-801 PYR MVL: 0.23 ± 0.01; *p* < 0.001; BS MK-801 INT MVL: 0.35 ± 0.01; MK-801 INT MVL: 0.3 ± 0.01, p < 0.001). However, NMDAr blockade did not modify the mean preferred phase of INTs (BS MK-801 INT Phase: 3.72 ± 0.09 rad; MK-801 INT Phase: 3.62 ± 0.11 *p* = 0.38),

Interestingly, in the vehicle condition, we observed an increase in the mean vector length (MVL) for PYR, while no changes were detected in their preferred phase (see Fig XXX), (BS Vehicle PYR MVL: 0.24 ± 0.01, Vehicle PYR MVL: 0.25 ± 0.01, p = 0.002; BS Vehicle PYR Phase: 3.79 ± 0.04 rad; Vehicle PYR Phase: 3.83 ± 0.04; p = 0.27). INTs showed no changes in their mean preferred phase (BS Vehicle INT Phase: 3.66 ± 0.09 rad; Vehicle INT Phase: 3.78 ± 0.09.; p = 0.39) or MVL (BS Vehicle INT MVL: 0.33 ± 0.01; Vehicle: 0.34 ± 0.01; p = 0.9).

Next, we investigated low (20-60 Hz) gamma oscillations. We found no significant changes on the preferred phase for both PYR and INT in the MK-801 condition (BS M-801 PYR Phase: 2.42 ± 0.04 rad; MK801 PYR Phase: 2.44 ± 0.06 rad; p = 0.34; MK-801 INT Phase: 2.37 ± 0.11 rad; MK-801 INT Phase: 2.49 ± 0.14 rad; p = 0.86). Moreover, NMDAr blockade reduced the strength of single neuron to gamma phase locking of INT, while no differences were found for PYR. (BS MK-801 PYR MVL: 0.13 ± 0.004; MK801 PYR MVL: 0.14 ± 0.005; p = 0.38; BS MK-801 INT MVL: 0.16 ± 0.01; MK-801 INT MVL: 0.12 ± 0.01; p = 0.004).

**Figure 3:**
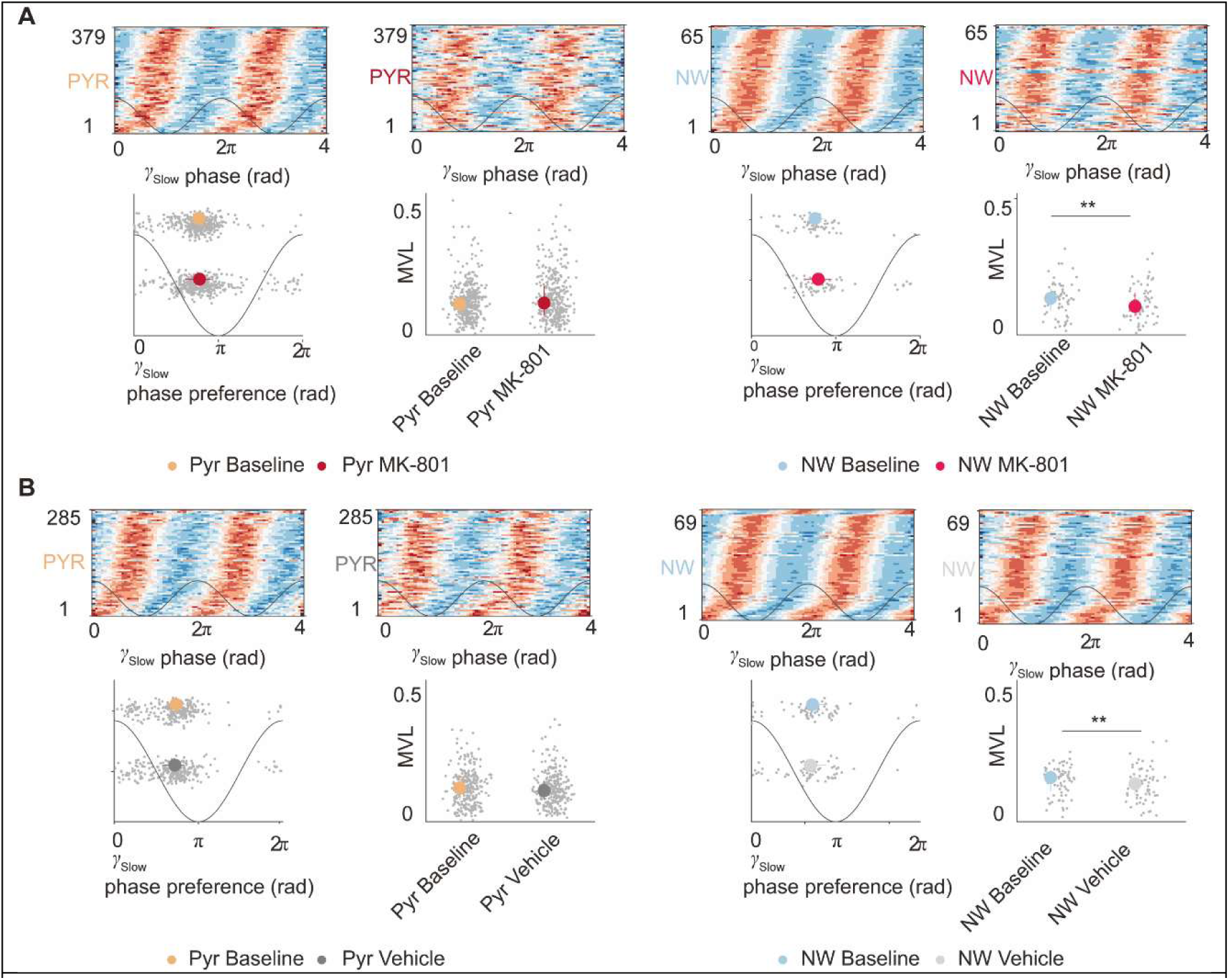
NMDAr blockade disrupts phase modulation of single neurons by slow gamma oscillations (γ slow). (A) Effects of MK-801 on phase modulation of pyramidal (PYR) and narrow waveform (NW) interneurons. Top row: Phase-locking of PYR (left) and NW (right) neurons to slow gamma (γ slow) oscillations during baseline and MK-801 conditions. Each heatmap represents the phase preference of neurons (y-axis) relative to the γ_slow phase (x-axis). Bottom row: Scatter plots showing phase preference of PYR and NW neurons (left) and modulation vector length (MVL) (right) in the baseline (gray/light blue) and MK-801 conditions (red). **MK-801 administration significantly reduced phase-locking strength (MVL) in NW interneurons. (B) Effects of vehicle administration on phase modulation of PYR and NW neurons. Top row: Phase-locking of PYR (left) and NW (right) neurons to γ_slow oscillations during baseline and vehicle conditions. Heatmaps show phase preference distribution. Bottom row: Scatter plots representing phase preference (left) and MVL (right) for PYR and NW neurons in the baseline (gray/light blue) and vehicle conditions (black). **No significant changes were observed in PYR neurons, while NW interneurons exhibited a significant reduction in MVL following vehicle administration (p < 0.01). These results indicate that NMDAr blockade with MK-801 disrupts phase modulation of NW interneurons by γ slow oscillations, whereas vehicle administration also affects NW modulation, suggesting a possible habituation or behavioral state effect.

In the vehicle condition we found that no significant differences in the phase of PYR and INT (BS Vehicle PYR Phase: 2.31 ± 0.05 rad; Vehicle PYR Phase: 2.27 ± 0.06 rad; p = 0.15; BS Vehicle INT Phase: 2.27 ± 0.14 rad; Vehicle INT Phase: 2.2 ± 0.11 rad; p = 0.39). PYR showed no changes in the MVL bua significant reduction of MVL in the vehicle condition for IN ((BS Vehicle PYR MVL: 0.14 ± 0.004; Vehicle PYR MVL: 0.13 ± 0.004; p = 0.59; BS Vehicle INT MVL: 0.19 ± 0.007; Vehicle INT MVL: 0.16 ± 0.008; p = 0.005).

**Figure 4.**
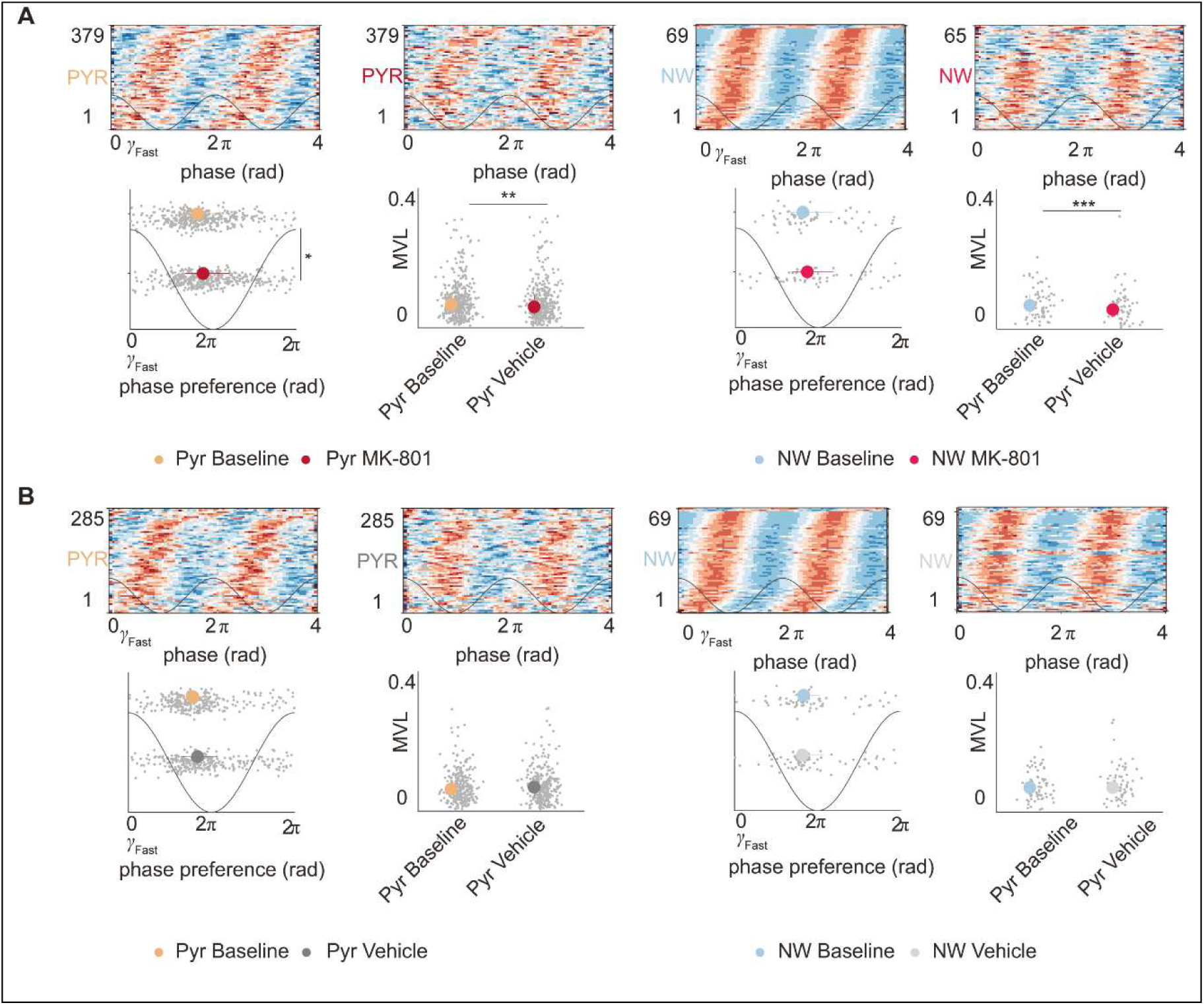
NMDAr blockade disrupts phase modulation of single neurons by fast gamma oscillations (γ fast). (A) Effects of MK-801 on phase modulation of pyramidal (PYR) and narrow waveform (NW) interneurons. Top row: Heatmaps representing the phase-locking of PYR (left) and NW (right) neurons to fast gamma (γ fast) oscillations during baseline and MK-801 conditions. The y-axis represents neuron index, while the x-axis shows the γ fast phase. Bottom row: Scatter plots illustrating phase preference (left) and modulation vector length (MVL) (right) for PYR and NW neurons during baseline (gray/light blue) and MK-801 (red) conditions. **MK-801 administration significantly reduced MVL in both PYR and NW interneurons, indicating weakened phase modulation. (B) Effects of vehicle administration on phase modulation of PYR and NW neurons. Top row: Heatmaps depicting the phase-locking of PYR (left) and NW (right) neurons to γ fast oscillations during baseline and vehicle conditions. Bottom row: Scatter plots showing phase preference (left) and MVL (right) for PYR and NW neurons in the baseline (gray/light blue) and vehicle (black) conditions. No significant changes were observed in PYR neurons, whereas NW interneurons exhibited a reduction in MVL following vehicle administration. These findings indicate that NMDAr blockade with MK-801 significantly impairs phase modulation of both PYR and NW neurons by γ fast oscillations, while vehicle administration has a minor effect, suggesting a potential influence of behavioral state changes.

Finally, we investigated the relationship between single neuron activity and high gamma oscillations. We found that NMDAr blockade significantly decreased the association strength between pyramidal neurons (PYR) and interneurons (INT) with high gamma (BS MK-801 PYR MVL: 0.06 ± 0.003; MK-801 PYR MVL: 0.05 ± 0.003; *p* = 0.005; BS MK-801 INT MVL: 0.07 ± 0.005; MK-801 INT MVL: 0.06 ± 0.006; *p* < 0.001). There was also a significant change in the mean preferred phase of PYR, but not of INT (BS MK-801 PYR Phase: 2.56 ± 0.06 rad; MK-801 PYR Phase: 2.76 ± 0.06; *p* = 0.02; BS MK-801 INT Phase: 2.55 ± 0.15 rad, MK-801 INT Phase: 2.73 ± 0.13, p = 0.11).

No effects were observed for any parameter during the vehicle condition for PYR cells, neither the mean preferred phase (BS Vehicle PYR Phase: 2.44 ± 0.07 rad; Vehicle PYR Phase: 2.62 ± 0.08 rad; *p* = 0.09) nor the MVL (BS Vehicle PYR MVL: 0.06 ± 0.003; Vehicle: 0.07 ± 0.003; *p* = 0.051) showed significant changes. Similarly, for INT cells, neither the preferred phase (BS Vehicle INT Phase: 2.62 ± 0.16 rad; Vehicle INT Phase: 2.57 ± 0.16; *p* = 0.69) nor the MVL (BS Vehicle INT MVL: 0.07 ± 0.005; Vehicle INT MVL: 0.07 ± 0.006, p = 0.54).

In summary, NMDAr blockade profoundly affected the modulation of PYR and INT activity by both theta and high gamma oscillations.

**Figure 5:**
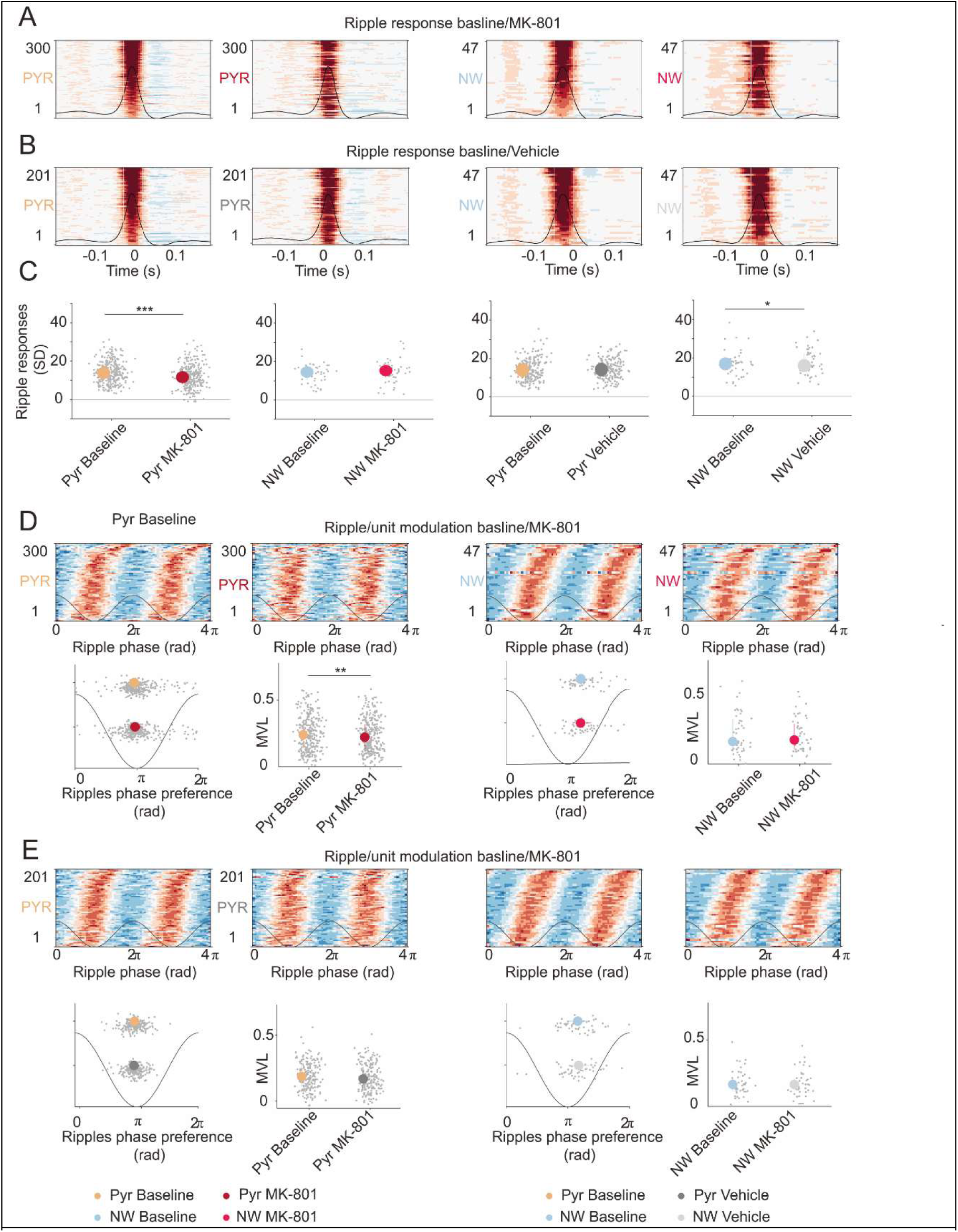
NMDAr blockade disrupts phase modulation of single neurons by fast gamma oscillations (γ fast). (A) Effects of MK-801 on phase modulation of pyramidal (PYR) and narrow waveform (NW) interneurons. Top row: Heatmaps representing the phase-locking of PYR (left) and NW (right) neurons to fast gamma (γ fast) oscillations during baseline and MK-801 conditions. The y-axis represents neuron index, while the x-axis shows the γ_fast phase. Bottom row: Scatter plots illustrating phase preference (left) and modulation vector length (MVL) (right) for PYR and NW neurons during baseline (gray/light blue) and MK-801 (red) conditions. **MK-801 administration significantly reduced MVL in both PYR (p < 0.05, marked by *) and NW interneurons (p < 0.001, marked by *), indicating weakened phase modulation. (B) Effects of vehicle administration on phase modulation of PYR and NW neurons. Top row: Heatmaps depicting the phase-locking of PYR (left) and NW (right) neurons to γ_fast oscillations during baseline and vehicle conditions. Bottom row: Scatter plots showing phase preference (left) and MVL (right) for PYR and NW neurons in the baseline (gray/light blue) and vehicle (black) conditions. No significant changes were observed in PYR neurons, whereas NW interneurons exhibited a reduction in MVL following vehicle administration. These findings indicate that NMDAr blockade with MK-801 significantly impairs phase modulation of both PYR and NW neurons by γ_fast oscillations, while vehicle administration has a minor effect, suggesting a potential influence of behavioral state changes.

### NMDAr blockade altered the dynamics of neuronal populations during periods of memory consolidation

Sharp-wave ripple oscillations (SWR; 120-200 Hz) are characteristic of resting, consummatory behaviors and are present in slow-wave sleep, contributing to consolidation (Diba & Buzsáki, 2007). This oscillatory activity depends on a precise excitatory and inhibitory balance that when is broken generates aberrant oscillations and memory distortion (Ibarz et al., 2010). As previously described, NMDAr blockade significantly change the spike train property of neurons altering the time organization of neurons activity. Thus, we expected to find changes in neurons activity during SWR. We investigated this by detecting SWR events and characterizing neuron response and modulation during these events.

NMDAr blockade generated a significant reduction in the recruitment of pyramidal cells during SWR (BS MK-801 PYR: 13.93 ± 0.33; MK-801 PYR: 11.65 ± 0.34; p < 0.001) while no significant changes were observed for narrow waveform interneurons (BS MK-801 INT: 14.51 ± 0.7; MK-801 INT: 15.28 ± 0.89; p = 0.88). In the vehicle condition, no changes were found for pyramidal neurons (BS Vehicle PYR: 13.99 ± 0.4; Vehicle PYR: 14.27 ± 0.34; p = 0.94), whereas narrow waveform interneurons showed a significantly decreased response during sharp-wave ripple events (BS Vehicle INT: 16.93 ± 0.94; Vehicle INT: 15.98 ± 0.96; p = 0.03).

We then analyzed the preferred phase and the strength of this association. In the MK-801 condition, we found no changes in the mean preferred phase (BS MK-801 PYR Phase: 3.01 ± 0.05 rad; MK-801 PYR Phase: 3.05 ± 0.05 rad; p = 0.51). However, in coherence with the reduced response during the MK-801 condition, we observed a significant reduction in the strength of the relationship between pyramidal neurons and SWR (BS MK-801 PYR MVL: 0.25 ± 0.01; MK-801 PYR MVL: 0.23 ± 0.01; p = 0.003). On the other hand, no significant changes were observed for narrow waveform interneurons, either in the mean preferred phase (BS MK-801 INT Phase: 3.8 ± 0.11; MK-801 INT Phase: 3.79 ± 0.13; p = 0.84) or the MVL (BS MK-801 PYR MVL: 0.17 ±; MK-801: 0.19 ± 0.02; p = 0.19).

In the case of the baseline vs vehicle condition we observed no changes either in the phase or the strength of the association for both PYR (BS Vehicle PYR Phase: 3.02 ± 0.03 rad; Vehicle PYR Phase: 3 ± 0.03; p = 0.62; BS Vehicle PYR MVL: 0.25 ± 0.01, Vehicle PYR MVL: 0.23 ± 0.01; p = 0.08) and INT (BS Vehicle INT Phase: 3.64 ± 0.13 rad; Vehicle INT Phase: 3.69 ± 0.14 rad; p = 0.62; BS Vehicle INT MVL: 0.17 ± 0.01, Vehicle INT MVL: 0.17 ± 0.01; p = 0.66).

**Figure 6:**
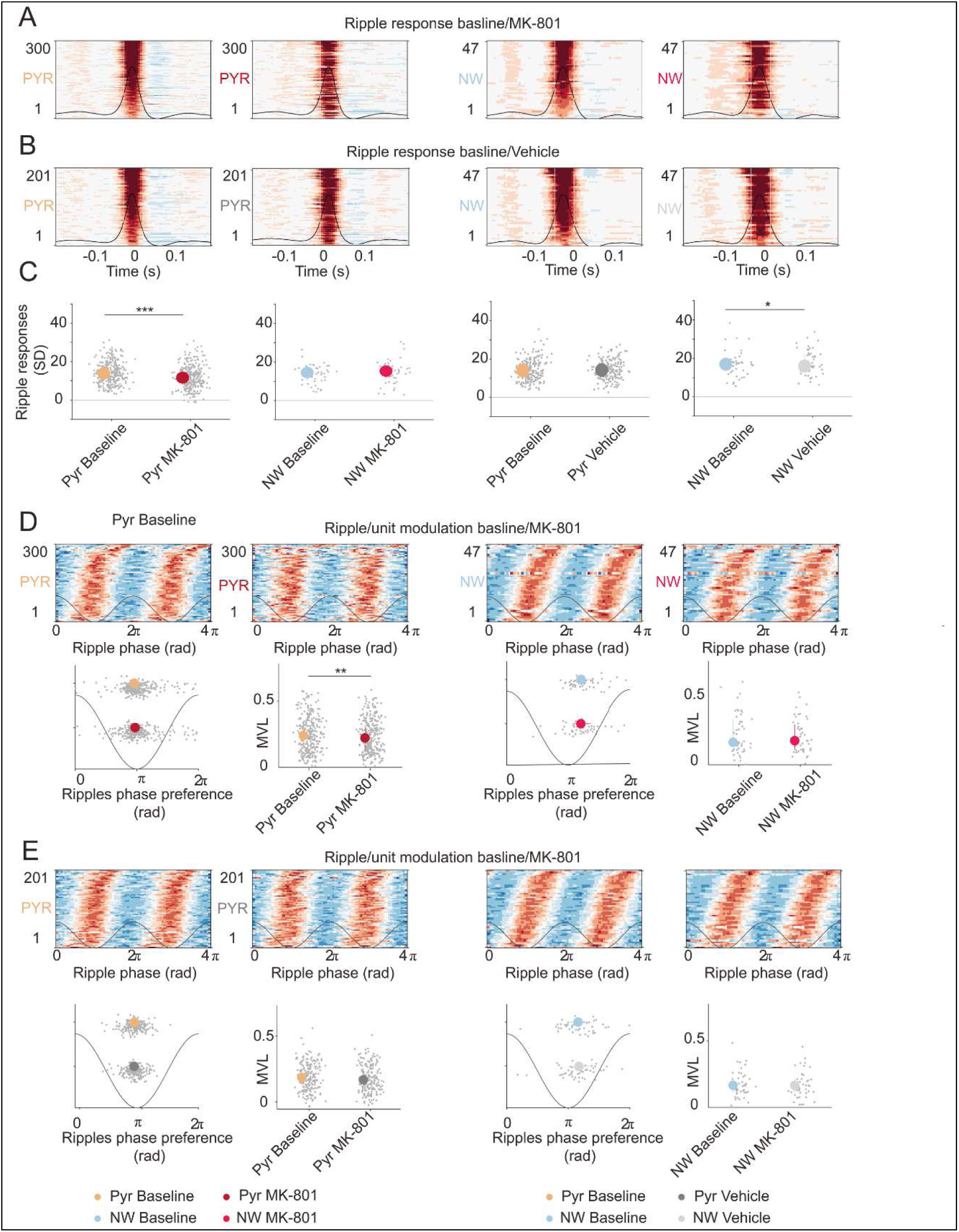
Effects of NMDAr blockade on ripple-related neuronal responses and phase modulation. (A) Ripple response in pyramidal (PYR) and narrow waveform (NW) interneurons under baseline and MK-801 conditions. Heatmaps showing peri-event time histograms (PETHs) of PYR (left) and NW (right) neurons aligned to ripple events (time 0) during baseline and MK-801 conditions. Warmer colors indicate increased firing probability. The black trace represents the average ripple waveform. (B) Ripple response in PYR and NW neurons under baseline and vehicle conditions. Similar to (A), showing PETHs of PYR and NW neurons aligned to ripple events during baseline and vehicle conditions. (C) Quantification of ripple-related responses. Scatter plots showing the strength of ripple responses (SD) in PYR and NW neurons for baseline vs. MK-801 (left) and baseline vs. vehicle (right). **Significant reductions in ripple responses were observed for PYR neurons following MK-801 administration for NW neurons following vehicle administration. (D) Ripple phase modulation of PYR and NW neurons in baseline vs. MK-801 conditions. Top row: Heatmaps showing spike-phase relationships of PYR and NW neurons with ripple oscillations under baseline and MK-801 conditions. Bottom row: Scatter plots illustrating phase preference (left) and modulation vector length (MVL) (right) for PYR and NW neurons. **MK-801 significantly reduced the phase modulation strength (MVL) of PYR neurons. (E) Ripple phase modulation of PYR and NW neurons in baseline vs. vehicle conditions. Top row: Heatmaps showing ripple phase-locking of PYR and NW neurons during baseline and vehicle conditions. Bottom row: Scatter plots displaying phase preference and MVL for PYR and NW neurons. No significant changes were observed in ripple phase modulation following vehicle administration. These results indicate that NMDAr blockade with MK-801 disrupts ripple-related firing and weakens phase modulation of pyramidal neurons, suggesting impaired hippocampal network function during memory consolidation.

In summary, NMDAr blockade altered the recruitment of pyramidal neurons, and the strength of the association to SWR, while no changes were observed in the case of the baseline vs vehicle condition.

**Figure 7:**
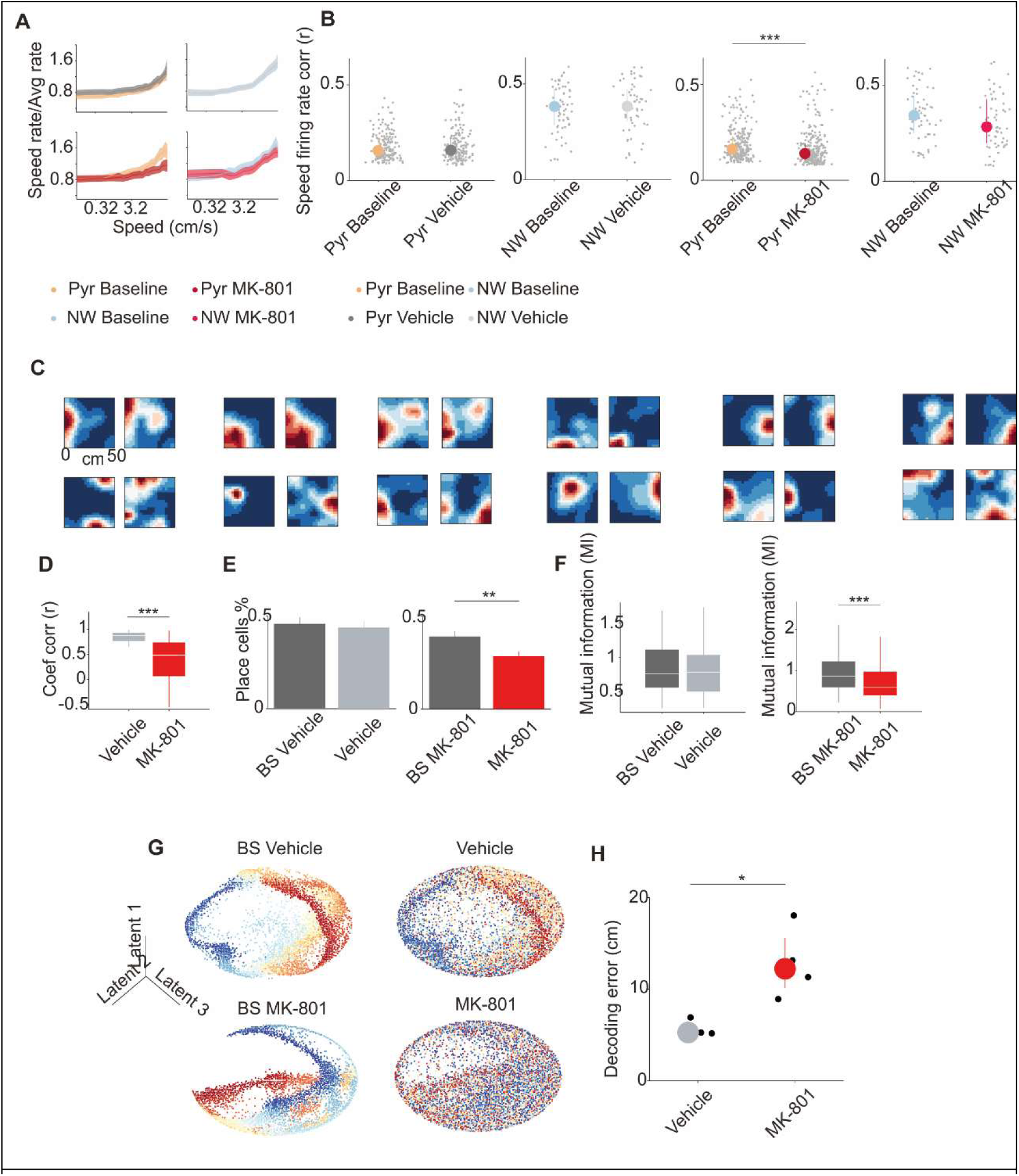
Effects of NMDAr blockade on hippocampal spatial coding and movement-related firing. (A) Relationship between firing rate and movement speed under different conditions. Plots show the average firing rate as a function of movement speed for pyramidal (PYR) and narrow waveform (NW) interneurons during baseline, vehicle, and MK-801 conditions. MK-801 administration disrupted the normal speed-firing relationship. (B) Correlation between firing rate and movement speed. Scatter plots depict the correlation coefficients (r) between firing rate and movement speed for PYR and NW neurons across baseline, vehicle, and MK-801 conditions. MK-801 significantly reduced speed coding in PYR neurons. (C) Place cell activity maps. Representative spatial firing rate maps of neurons recorded in the open field under different conditions. Warmer colors indicate higher firing rates. MK-801 treatment impaired place cell stability and spatial representation. (D) Spatial correlation of place fields. Box plots comparing the correlation of spatial firing patterns between baseline and drug conditions. MK-801 significantly reduced spatial correlation. (E) Percentage of place cells. Bar plots comparing the proportion of neurons classified as place cells across conditions. MK-801 significantly reduced the number of place cells compared to baseline. (F) Mutual information analysis. Box plots showing the mutual information (MI) between neuronal firing and spatial location. MK-801 significantly reduced MI values compared to baseline. (G) Latent space representations of neural activity. 2D projections of neural population activity under different conditions using dimensionality reduction. MK-801 altered the neural state-space organization compared to vehicle and baseline conditions. (H) Decoding error analysis. Scatter plot showing the decoding error of spatial position based on neural activity. MK-801 significantly increased the decoding error, indicating impaired spatial representation. These results demonstrate that NMDAr blockade with MK-801 disrupts hippocampal spatial coding, weakens movement-related firing rate modulation, and impairs the ability to accurately represent spatial information, which may underlie cognitive deficits observed in schizophrenia

### NMDAr blockade induced hyperlocomotion and disrupted the correlation between the firing rate of pyramidal cells in the hippocampus and the speed of movement

We found a significantly increased exploration in the open field after NMDAr blockade, as measured by the distance traveled per minute (BS: 2.87 ± 0.3; MK801: 6.09 ± 1.08; p = 0.04) and the mean speed of movement (BS: 5.62 ± 0.62; MK801: 12.13 ± 1.61; p = 0.04). On the contrary, and as expected from previous results, we found significantly reduced exploration in the vehicle condition for both the distance traveled per minute (BS: 3.03 ± 0.43; Vehicle: 1.40 ± 0.49; p < 0.03) and the mean speed of movement (BS: 5.08 ± 0.47; Vehicle: 3.78 ± 0.74; p < 0.03). These results indicate a behavioral effect of our pharmacological manipulation but also suggest habituation in the case of the vehicle condition.

In our previous work, we observed a rupture in the physiological association of oscillatory activity and the speed of movement(Abad-Perez et al., 2023). We aimed to understand if there was a similar effect in the correlation of the firing rate of single neurons and the speed of movement after NMDAr blockade. We computed the instantaneous firing rate of CA1 neurons in each position sample (3 ms bin) and calculated the Pearson correlation between the instantaneous firing rate and the speed of movement, obtaining a speed score correlation coefficient and an associated p-value. Only significantly positively correlated cells for both baseline and drug conditions were analyzed, and epochs of quietness were not included in the analysis. We observed a reduced speed score coefficient in pyramidal cells after NMDAr blockade (BS: 0.10 ± 0.01; MK801: 0.08 ± 0.01; p < 0.001), while no differences were found for interneurons (BS: 0.33 ± 0.02; MK801: 0.26 ± 0.02; p = 0.12). In the vehicle condition, we found no changes for either pyramidal cells (BS: 0.1 ± 0.01; Vehicle: 0.1 ± 0.01; p = 0.69) or interneurons (BS: 0.38 ± 0.02; Vehicle: 0.38 ± 0.02; p = 0.51).

### NMDAr blockade induced a distortion of the spatial signal

We found that NMDAr blockade significantly impaired spatial representation in the hippocampus. This was reflected by a reduction in the correlation of firing rate maps between baseline and drug conditions (Vehicle: 0.88 ± 0.02; MK-801: 0.48 ± 0.04). Similarly, the percentage of neurons classified as place cells was lower after MK-801 administration (BS MK-801: 0.38 ± 0.48; MK-801: 0.28 ± 0.45), whereas no significant changes were observed in the vehicle condition (BS Vehicle: 0.49 ± 0.5; Vehicle: 0.47 ± 0.5).

Additionally, mutual information between neuronal firing and spatial location was significantly reduced after MK-801 administration (BS-MK801: 0.88 ± 0.05; MK-801: 0.61 ± 0.04), whereas no significant changes were observed in the vehicle condition (BS Vehicle: 0.8 ± 0.05; Vehicle: 0.78 ± 0.05). Finally, decoding error, a measure of spatial representation accuracy, was significantly increased in the MK-801 condition (Vehicle: 5.26 ± 0.56; MK-801: 12.23 ± 1.93), suggesting that NMDAr blockade impaired the ability of hippocampal neurons to encode spatial information accurately.

These results indicate that NMDAr hypofunction disrupts spatial coding in the hippocampus, leading to a loss of place cell stability and a reduced ability to represent spatial information, which may contribute to cognitive deficits associated with schizophrenia models.

## Discussion

Our findings delineate the extensive impact of NMDAr blockade on the activity of CA1 pyramidal neurons and interneurons. These effects encompass alterations in spike train organization, and modifications in the coupling strength between oscillatory patterns and spike timing across different frequency bands. Additionally, a diminished correlation between neuronal firing rates and locomotor velocity was observed and a significant alteration of the spatial signal. Moreover, we found similar alterations in gamma activity and theta/gamma co-modulation as previously described (Abad-Perez et al., 2023). These outcomes are consistent with the reported changes in oscillatory dynamics and cognitive impairments documented in prior literature, including our own investigations (Abad-Perez et al., 2023; Bygrave et al., 2019).

### Firing rate and spike train desynchronization indicates excitatory-inhibitory imbalance after NMDAr blockade

The integrity of physiological oscillatory activity critically relies on the appropriate balance between excitation and inhibition. NMDAr blockade possesses the potential to disrupt this balance, leading to oscillopathies and cognitive deficits reminiscent of those observed in patients with schizophrenia (Uhlhaas & Singer, 2010). Our findings revealed that MK-801 significantly impacted this excitation-inhibition balance, resulting in an alteration in the spike trains of both pyramidal neurons and interneurons. Additionally, we observed a reduction in the bursting properties of pyramidal neurons, suggesting a disruption in the temporal organization of spike trains. Such alteration may represent a mechanism underlying impaired information transmission between brain regions (Kepecs et al., 2002; Lisman & Jensen, 2013). Similar observations have been informed in the prefrontal cortex (Jackson et al., 2004; Molina et al., 2014), although reports also suggest differential effects on pyramidal neurons and interneurons (Homayoun & Moghaddam, 2007). Specifically, an increase in the firing rate of pyramidal neurons in the prefrontal cortex was noted, seemingly driven by an initial reduction in the firing rate of interneurons. However, we did not observe similar temporal dynamics in the firing rates of pyramidal neurons and interneurons as described in the prefrontal cortex. These disparities may be attributed to differences in the circuit organization between the hippocampus and prefrontal cortex. Furthermore, these variations could underly the specific alterations in oscillatory activity induced by MK-801 across different brain regions (Abad-Perez et al., 2023).

In contrast to the effects obtained after NMDAr blockade, the firing rate of both groups of neurons was reduced in vehicle condition, possibly due to a habituation to the open field during the different conditions and the consequent reduction in the level of exploratory behavior(Abad-Perez et al., 2023).

Moreover, NMDAr blockade reduced the correlation between the firing rate of neurons and the and the speed of movement of both pyramidal cells and narrow waveform interneurons. This indicates a reduced sensitivity of the neuronal code to establish the speed of movement, and therefore a loss in spatial coding efficiency. Moreover, this same effect might be even more critical to speed-modulated cells located in the MEC that has been hypothesized to support firing of both place cells and grid cells by sending to them information about locomotion speed required for path integration ((Góis & Tort, 2018; Iwase et al., 2020)).

### NMDAr blockade disrupted timing and the modulation of neurons by oscillatory activity

As we expected, NMDAr blockade altered the modulation of neurons by different oscillatory activity, probing the disruption of network-unit interaction. We observed a reduction in the strength of modulation of pyramidal neurons during theta. Theta activity is thought to contribute to the encoding of information during exploration, and to organize the time firing of different population of neurons (Inostroza et al., 2013; Klausberger & Somogyi, 2008; Lopez-Pigozzi et al., 2016). Thus, this reduction in the modulation of pyramidal neurons by theta, as well as a reduction in their bursting properties, could be associated with an impairment in the computational capabilities of place cells and spatial cognition.

A major readout of NMDAr disfunction is the alteration of gamma oscillations ((Gonzalez-Burgos et al., 2023). We observed that NMDAr blockade strongly altered single neuron modulation by low and high gamma band, however finding some specificities.

In the case of the low gamma, pyramidal and interneurons advanced their preferred firing preference after the administration of MK-801. Moreover, pyramidal neurons showed a reduction in the strength of modulation in both, high and low gamma. These results indicate an apparent stronger impact of NMDAr blockade in the low gamma band. Low gamma activity has been associated with inputs from the CA3 to the stratum radiatum (SR) and therefore, with the storage of information, but also with successful spatial working memory (Bieri et al., 2014; Montgomery & Buzsáki, 2007). On the other hand, the high gamma band is related to inputs from medial entorhinal cortex, layer III, to the “stratum lacunosum moleculare” (SLM). It is though that these oscillations regulate the integration of spatial signals from the MEC (Valero et al, XXX). Alterations in the phase and the strength of modulation here described are coherent with a reduction in the computation capabilities of the hippocampal network to process spatial information and other memory processes.

Remarkably, we observed that the relationship between low gamma oscillations and pyramidal and interneurons during the vehicle condition was strengthened. This augmentation might reflect a higher level of coordination between the CA3 and CA1 region, probably reflecting the re-exposure of mice to the same environment. This could be an indicator of a memory trace of the environment. This idea is supported by previous results ((Fernández-Ruiz et al., 2017; Tort et al., 2009), and follow the same trend of our own results, demonstrating the increase of theta/gamma across different temporal epochs of the baseline and vehicle conditions that we observed in our previous work (Abad-Perez et al., 2023).

### Reduced neuronal population synchrony during NMDAr Blockade could relate to cognitive deficits

As previously mentioned, the synchronization between neuronal populations is one possible mechanism to generate functionally coherent neuronal supporting cognition. The reduction of this synchrony might underly cognitive symptoms in different disorders, including SCZ (Uhlhaas & Singer, 2010). We observed that NMDAr blockade reduce the population synchronization of pyramidal neurons during spontaneous behavior. This reduction in the coordination of neuronal activity might be related the changes in the bursting properties of pyramidal neurons, as well as in the increase in the firing rate.

### Impaired ripple response after NMDAr blockade might underly memory deficits

It is well established that ripple activity facilitates consolidation of new memories by enabling neuronal ensemble replay during slow wave sleep ((Jadhav et al., 2012; Kudrimoti et al., 1999). Moreover, disruption of artificial ripple enhancement produces memory impairment or potentiation respectively (Fernández-Ruiz et al., 2019; Girardeau et al., 2009). On the other hand, NMDAr blockade impairs new memory formation and place cells stabilization, possibly by altering the balance between synaptic potentiation and depression during the learning process (Dupret et al., 2010; Kentros et al., 1998)We observed that NMDAr blockade decrease the recruitment of pyramidal neurons during ripples, and the strength of ripple to unit modulation. Thus, our results indicate a disfunction in SWR activity of pyramidal neurons might underly the consolidation of neurons by altering spike time dependent plasticity, and LTP-like mechanisms supporting memory consolidation (Lüscher & Malenka, 2012) (Lüscher et al, 2012). On the hand, neurons showed an enhanced response during the vehicle condition. This might be due to an increase in the consolidation process.

**In summary,** the NMDAr blockade generated the alteration of the E-I balance and spike train, and the impairment in the probability of information transmission across brain regions. This alteration might cause a functional disconnection between brain regions and might explain multiple cognitive deficits observed in animal models of NMDAr hypofunction and in SCZ patients. All these elements can explain the distortion of place cell coding and the diminished correlation between firing rate and the speed of movement.

As previously mentioned, oscillatory activity provides specific time windows for different types of neurons to be active (Klausberger and Somogy, 2008). The observed distortion here indicates the existence of widespread changes in spike time activity, and therefore the potentiality to impair cognitive processing by hippocampal circuits. This alteration could cascade to other brain regions such as the PFC, altering the communication between regions and therefore cognitive processing, for example during spatial working memory. Moreover, these changes could explain oscillatory changes and the co-modulation of different brain rhythms, for example theta/gamma co-modulation disruption during working memory.

## Contributions

JRBM, PA and RR designed the experiments. JRBM, PA and GR implemented the experiments. PA developed the analysis toolbox. JRBM, PA, FJMP, GR and ATZ participated in the analysis of the data. GC processed the tissue for anatomical analysis. JRBM and PA discussed and interpreted the results. VB, LM, AF supported the project providing resources for its development. JRBM, PA, MV, VB, LM, GR, FJMP, ATZ and AF discussed the results. JBRM wrote the paper.

## Acknowledgments

We want to acknowledge the members of Victor Borrell Lab, Luís Marinez Otero for their help and support. In addition, we want to thank Manuel Valero and his team for their support.

## Funding

This project was supported by the Spanish State Research Agency, ‘‘Ministerio de Ciencia, Innovación y Universidades” (RTI2018-097474-A-100) obtained by JRBM.

JRBM was supported by Ministerio de Ciencia, Innovación y Universidades” (RTI2018-097474-A-100), PA was funded by the UCHCEU-Banco de Santander fellowship granted to PA and JRBM and the internship funding scheme of UCH-CEU.

“Effect of RO compounds on the electrophysiological coupling of hippocampal-prefrontal circuits”. F. Hoffmann-La Roche Ltd. by obtained by JRBM.

VB was supported by grants from the European Research Council (309633) to Victor Borrell, Spain and the Spanish State Research Agency (PGC2018-102172-B-I00, as well as through the “Severo Ochoa” Programme for Centers of Excellence in R&D, ref. SEV-2017-0723).

LM was supported by ERC-2020-SyG 951631 – XSCAPE Project on Material Minds.

